# Hypermethylation of human DNA: Fine-tuning transcription associated with development

**DOI:** 10.1101/212191

**Authors:** Carl Baribault, Kenneth C. Ehrlich, V. K. Chaithanya Ponnaluri, Sriharsa Pradhan, Michelle Lacey, Melanie Ehrlich

**Affiliations:** Tulane Cancer Center, Tulane University Health Sciences Center, New Orleans, LA 70112, USA.; Department of Mathematics, Tulane University, New Orleans, LA 70118, USA; Center for Bioinformatics and Genomics, Tulane University Health Sciences Center; New England Biolabs, Ipswich, MA 01938, USA; Hayward Genetics Center Tulane University Health Sciences Center, New Orleans, LA 70112, USA.

**Keywords:** DNA methylation, chromatin, development, epigenetic memory, CTCF, *NR2F2*(COUP-TFII), *NKX2-5*, *LXN* (Latexin), *EN1*, *PAX3*

## Abstract

Tissue-specific gene transcription can be affected by DNA methylation in ways that are difficult to discern from studies focused on genome-wide analyses of differentially methylated regions (DMRs). We studied 95 genes in detail using available epigenetic and transcription databases to detect and elucidate less obvious associations between development-linked hypermethylated DMRs in myoblasts (Mb) and cell-and tissue-specific expression. Many of these genes encode developmental transcription factors and display DNA hypermethylation also in skeletal muscle (SkM) and a few heterologous samples (e.g., aorta, mammary epithelial cells, or brain) among the 38 types of human cell cultures or tissues examined. Most of the DMRs overlapped transcription regulatory elements, including canonical, alternative, or cryptic promoters; enhancers; CTCF binding sites; and long-noncoding RNA (lncRNA) gene regions. Among the prominent relationships between DMRs and expression was promoter-region hypermethylation accompanying repression in Mb but not in many other repressed samples (26 genes). Another surprising relationship was down-modulated (but not silenced) expression in Mb associated with DNA hypermethylation at cryptic enhancers in Mb although such methylation was absent in both non-expressing samples and highly expressing samples (24 genes). The tissue-specificity of DNA hypermethylation can be explained for many of the genes by their roles in prenatal development or by the tissue-specific expression of neighboring genes. Besides elucidating developmental epigenetics, our study provides insights into the roles of abnormal DNA methylation in disease, e.g., cancer, Duchenne muscular dystrophy, and congenital heart malformations.

## Introduction

Development-related changes in DNA methylation play various roles in controlling expression of differentiation-related genes in mammals ^1–5^. Still, much remains to be learned about the functional significance of tissue-and development-specific differential DNA methylation. In vitro methylation throughout CpG-rich active promoters usually leads to silencing of the associated genes, and, during development, some promoters are repressed in conjunction with DNA hypermethylation ^6–8^. Recent findings about the prevalence of unstable antisense (AS) transcripts at active promoter regions ^9^ and long non-coding RNA (lncRNA) genes near promoters ^10, 11^ demonstrate the need for more investigation of the roles that DNA methylation plays in modulating development-linked gene expression from the vicinity of promoters.

The effects of gene-body DNA methylation on gene expression are known to be complex ^6^ and have been confusing ^12^, especially in view of intragenic enhancers, cryptic promoters, and alternative promoters ^1, 13, 14^. Gene-body methylation is positively associated with gene expression in some genome-wide studies ^15, 16^ but also has been reported to have a negative association with transcription ^3^. In addition, intragenic DNA methylation has been implicated in helping regulate the choice of exon-intron boundaries during co-transcriptional splicing of pre-mRNAs ^17^. Both intragenic and Intergenic undermethylated enhancers ^18^, which are often bidirectionally transcribed to give short, transient enhancer RNAs (eRNAs) ^9^, are critical in development-associated transcription control ^18^. Confounding the analysis of the biological effects of differential DNA methylation on expression is the finding that many changes in DNA methylation during development or oncogenesis do not correlate with changes in expression of the associated gene ^3, 8, 19^.

While whole-genome studies have elucidated many important relationships between development-linked epigenetics and transcription ^3,15, 16, 20–24^, there are some complicated, modest, or infrequent associations that they may miss. We investigated the associations between myogenic differentially methylated regions (DMRs) at or near 95 genes and within their surrounding gene neighborhoods. In this study, untransformed human muscle progenitor cells, myoblasts (Mb) and myotubes (Mt), as well as skeletal muscle tissue (SkM) were compared with many dissimilar cell cultures or tissues. We focused on the SkM lineage because SkM normally contributes the most mass to the human body; plays a vital and dynamic role in many disparate bodily functions; is involved in many congenital and somatic diseases (including cancer); is subject to frequent postnatal repair; and has a major role in aging ^25–27^. The genome-wide epigenetics specific to this dynamic tissue and to Mb and Mt are beginning to be studied in detail ^11, 26, 28, 29^. The importance of DNA methylation in myogenesis was demonstrated by conditional knockout of one of the three DNA methyltransferase genes ^30^, *Dnmt3a*, in the mouse SkM lineage ^31^. Moreover, postnatal promoter methylation changes in SkM have been implicated in muscle physiology as exemplified by a rat model of atrophy-disuse ^32^. Our study elucidates how DNA methylation can fine-tune gene expression in normal human development, not only in the SkM lineage, but also in surprisingly diverse cell lineages that shared DNA hypermethylation with myogenic cells in some gene regions or that were distinguished from myogenic cells by opposite methylation patterns at other gene regions.

## Results

### Genome-wide overview of relationships between myogenic DNA hypermethylation and chromatin structure

Before focusing on individual genes, we examined genome-wide relationships between myogenic differential DNA methylation and chromatin structure. For these comparisons, we used ENCODE-derived chromatin state segmentation profiles ^21^ and our previously described database of almost 10000 CpG sites that are significantly hypermethylated and a similar number that are hypomethylated in myogenic vs. non-muscle cell cultures (Figure 1). Significantly differentially methylated sites (DM sites) in myogenic progenitor cells (the set of Mb and Mt) vs. 16 non-myogenic cell cultures were determined by reduced representation bisulfite sequencing (RRBS) ^20, 28^. For simplicity we refer to Mb and Mt DM sites (which were very similar in DNA methylation) as Mb DM sites. The highest ratio of Mb-hypermethylated to Mb-hypomethylated sites was found at chromatin with the histone marks of actively transcribed gene regions (txn-chromatin, enriched in H3 lysine-36 trimethylation, H3K36me3; Figure 1a). Mb-hypermethylated sites embedded in txn-chromatin were seen not only in the bodies of genes but also upstream of the transcription start site (TSS) and especially in intergenic regions (Figure 1b). The lowest ratio of Mb-hypermethylated to Mb-hypomethylated sites was seen in strong enhancer chromatin (enh-chromatin; enriched in H3K4me1 and H3 lysine-27 acetylation, H3K27ac) in both intragenic and intergenic regions (Figure 1a-c). Weak enh-chromatin (H3K4me1 with low or no H3K27ac) was also enriched in Mb-hypomethylated vs. Mb-hypermethylated sites but not as much as for strong enh-chromatin (Figure 1a). Mb hypermethylation generally favored weak over strong enh-chromatin irrespective of the gene subregion (Figure 1d and e) while the opposite was true for intragenic Mb hypomethylation (Figure 1d and e). Mb hypermethylation was seen preferentially in weak promoter chromatin (weak prom-chromatin; enriched in H3K4me3 but with only low H3K27ac) relative to strong prom-chromatin (H3K4me3 with strong H3K27ac signal; Figure 1a and Supplementary Figure S1a). These results suggest that DNA methylation in cis often fine-tunes, rather than just negates, enhancer or promoter activity, a conclusion supported by our analysis of individual genes described below.

**Figure 1.**
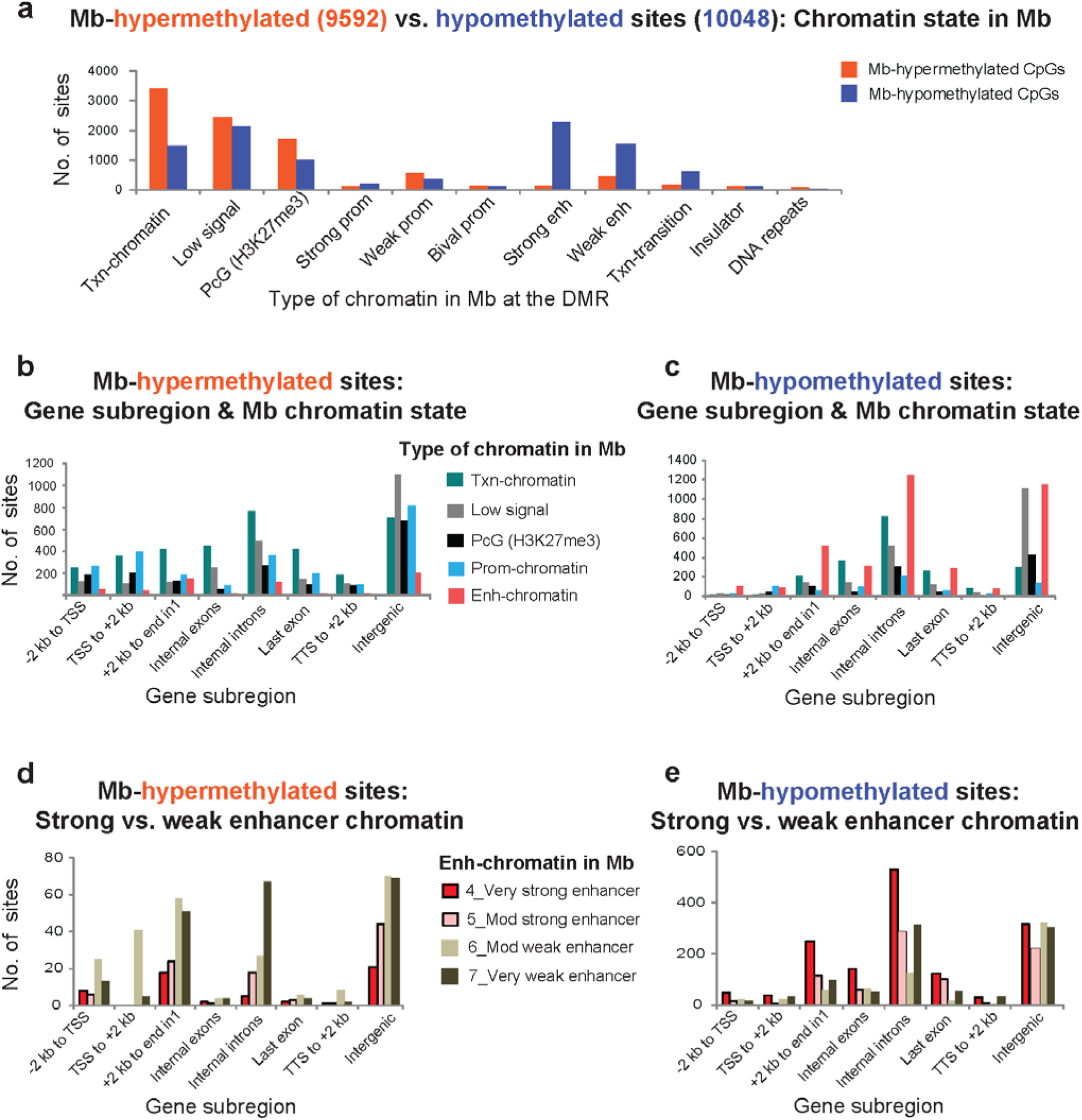
Mb-hypermethylated sites are particularly prevalent in transcription-type (txn) chromatin and Mb-hypomethylated sites in strong enhancer-type chromatin. **(a)** The 9592 CpGs that are significantly hypermethylated and the 10048 CpGs that are significantly hypomethylated in the set of Mb and Mt vs. 16 types of non-myogenic cell cultures were mapped to their chromatin state (15-state Chromatin State Segmentation by HMM from ENCODE/Broad ^21, 34^). **(b), (c)** The Mb-hypermethylated and Mb-hypomethylated sites are compared as to both their chromatin state and gene-subregion. Prom-chromatin, strong, weak or bivalent promoter chromatin; enh-chromatin, strong or weak enhancer chromatin; Low signal, little or no H3K4 methylation, H3K27me3, H3K27ac, or H3K36me3; TTS, transcription termination site. **(d), (e)** Distribution of Mb-hypermethylated and Mb-hypomethylated sites among very strong, moderately strong, moderately weak or very weak enhancer chromatin according to the 15-state chromatin segmentation ^21^.

We examined the chromatin segmentation states in Mb and in six non-myogenic cell cultures at Mb DM sites. For the Mb-hypermethylated sites, a wide variety of underlying chromatin states was seen in the non-Mb cell cultures with two exceptions (Supplementary Figure S1b). First, Mb-hypermethylated sites located in chromatin enriched in repressive H3K27me3 (PcG-chromatin) in Mb were usually also located in PcG-chromatin in the other cell types (Supplementary Figure S1b, dotted box) or, for H1 embryonic stem cells (ESC), in bivalent promoter chromatin (both repression-associated H3K27me3 and activation-associated H3K4me3; Supplementary Figure S1b, orange arrow). Second, whatever the chromatin state in Mb at the Mb-hypermethylated sites, the predominant chromatin state in ESC at the Mb-hypermethylated DM sites was bivalent prom chromatin (Figure 2b; Supplementary Figure S1b, arrows). Of the total Mb-hypermethylated sites, 59% overlapped ESC bivalent chromatin. Eighty-two percent of these Mb-hypermethylated sites at ESC bivalent chromatin regions were located in CpG islands (CGIs) while only 66% of all Mb-hypermethylated sites were at CGIs (Figure 2b), a significant enrichment in the former sites in CGI (p<0.0001). Therefore, the strong enrichment of ESC bivalent chromatin at Mb-hypermethylated sites can be explained largely by the overrepresentation of CGIs at the Mb-hypermethylated sites.

**Figure 2.**
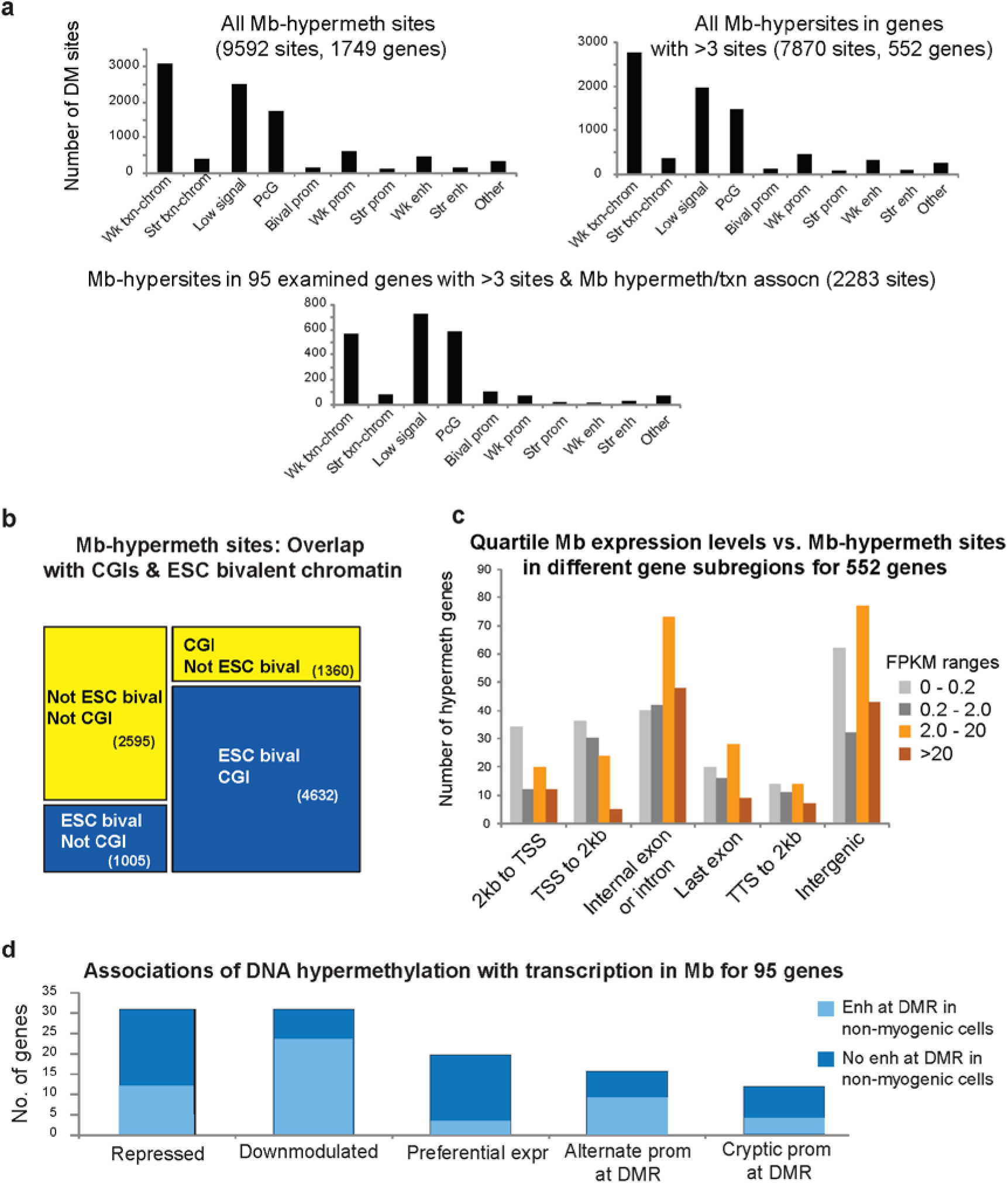
Mb-hypermethylated sites are strongly overrepresented in subregions displaying bivalent chromatin in ESC and have diverse associations with transcription. **(a)** There are similar distributions of Mb-hypermethylated sites among the different chromatin segmentation states (Figure 1a) whether all 9592 sites, the subset of 7870 sites that occur in clusters of at least three sites, or just the 2283 sites in the 95 genes highlighted in this study are considered. (b) Mb-hypermethylated sites are enriched in sites that overlap bivalent chromatin ^21^ in ESC, which can be largely explained by their overlap with CpG islands ^34^. (c) Distribution of Mb-hypermethylated sites among different Mb transcription classes by quartiles from RNA-seq profiles of Mb that used > 200 nt poly(A)^+^ RNA from ENCODE/Wold Lab at Caltech ^28, 34^ for genes associated with >3 Mb-hypermethylated sites. (d) The 95 genes associated with Mb-hypermethylated sites and selected for most of the analyses in this study fell into the indicated five major categories for their relationship of Mb hypermethylation with transcription in Mb as described in the text. These genes are individually annotated in Supplementary Tables S1 – S4. Light blue, the gene was associated with a Mb-hypermethylated DMR that displayed enhancer chromatin and a lack of DNA methylation in at least one non-myogenic cell type or tissue.

### Associations of myogenic hypermethylation with cell type-specific differences in chromatin state and expression

We next focused on relationships between myogenic hypermethylation and tissue or cell (T/C)-specific gene expression using Mb-hypermethylated sites mapped to the nearest gene ^28^ and RNA-seq-determined expression levels categorized as Mb FPKM divided into quartiles. Other than in the region from the TSS to 2 kb downstream (TSS to 2kb), the correlations between DNA hypermethylation and expression in Mb were not consistent (Figure 2c). Therefore, to determine if many biologically relevant correlations could be made we examined in detail the epigenetic and transcription parameters for many individual genes.

Using RRBS methylome and RNA-seq data for 9-18 types of cell cultures or the more comprehensive bisulfite-seq (BS-seq) methylome profiles for 20 types of tissues as well as chromatin epigenetic profiles for these cell cultures and tissues ^23, 28, 33, 34^, we screened 280 of the 552 genes (exclusive of multigenically regulated *HOX* cluster genes ^35^) that were associated with four or more Mb-hypermethylated sites to identify genes with T/C-specific relationships between hypermethylation and gene expression. From these we were able to choose 95 genes that displayed such positive or negative relationships (Supplementary Tables S1-S4). The distribution of hypermethylated sites over gene subregions for this gene subset was similar to that of the larger 552-and 1749-gene sets associated with RRBS-determined Mb-hypermethylated sites (Figure 2a). A whole-genome analysis showed very strong overrepresentation of the gene ontology terms “homeobox gene” and “sequence-specific DNA binding protein” ^28^. Similarly, 48 and 22 of the 95 genes that are the focus of the present study are associated with developmental TFs or other developmental proteins, respectively (Supplementary Tables S1a – S4a). Although we are using steady-state RNA levels (RNA-seq) as a marker of differential transcription, chromatin state segmentation profiles were consistent with the RNA-seq data. This indicates that post-transcriptional control of RNA levels was not interfering with our analysis.

In the set of 95 genes the two most frequently observed DNA methylation/transcription relationships were Mb hypermethylation being associated with repression in Mb (31 genes) or selective transcription in Mb but with expression at down-modulated levels relative to one or more other types of samples (31 genes; Figure 2d, Supplementary Tables S1 and S2). We refer to these as Mb-hypermeth/repr and Mb-hypermeth/downmod genes, respectively. *LXN, NKX2-5*, *SIX3*, *ZIC4*, *DBX1* and *LAD1* (Figure 3 and 4; Supplementary Figures S2 – S5) illustrate Mb-hypermeth/repr genes, and *NR2F2, ZIC1, PITX1, SIM1*, and *TBX3* (Figure 5, Supplementary Figures S3, S6 – S8) are examples of Mb-hypermeth/downmod genes. Other genes (21; Supplementary Table S3) displayed myogenic hypermethylation positively correlated with gene expression but with no examples of unmethylated samples having higher expression (Mb-hypermeth/pref-expr genes), e.g., *EN1*, *SIX2, LBX1* and *LBX1-AS1* (Figure 6 and Supplementary Figures S2 and S9). Some of the Mb-hypermeth/pref-expr genes might belong in the down-modulated category if there are specialized prenatal or postnatal cell types that have not yet been described as lacking methylation at the DMR and having higher expression of the associated gene than in the examined adult-derived Mb samples. An additional 29 genes were associated with DMRs overlapping alternative promoters (encoding a known or deduced RNA isoform derived from many of the canonical exons) or cryptic promoters (with a probable lncRNA isoform not annotated in RefSeq or Ensembl) in unmethylated samples (Supplementary Table S4). Among these genes were *ZIC1, DBX1, LAD1, LBX1*, and *RFX1* (Supplementary Figures S3-S5, S9, and S11). About half of these genes were also in one of the aforementioned categories.

**Figure 3.**
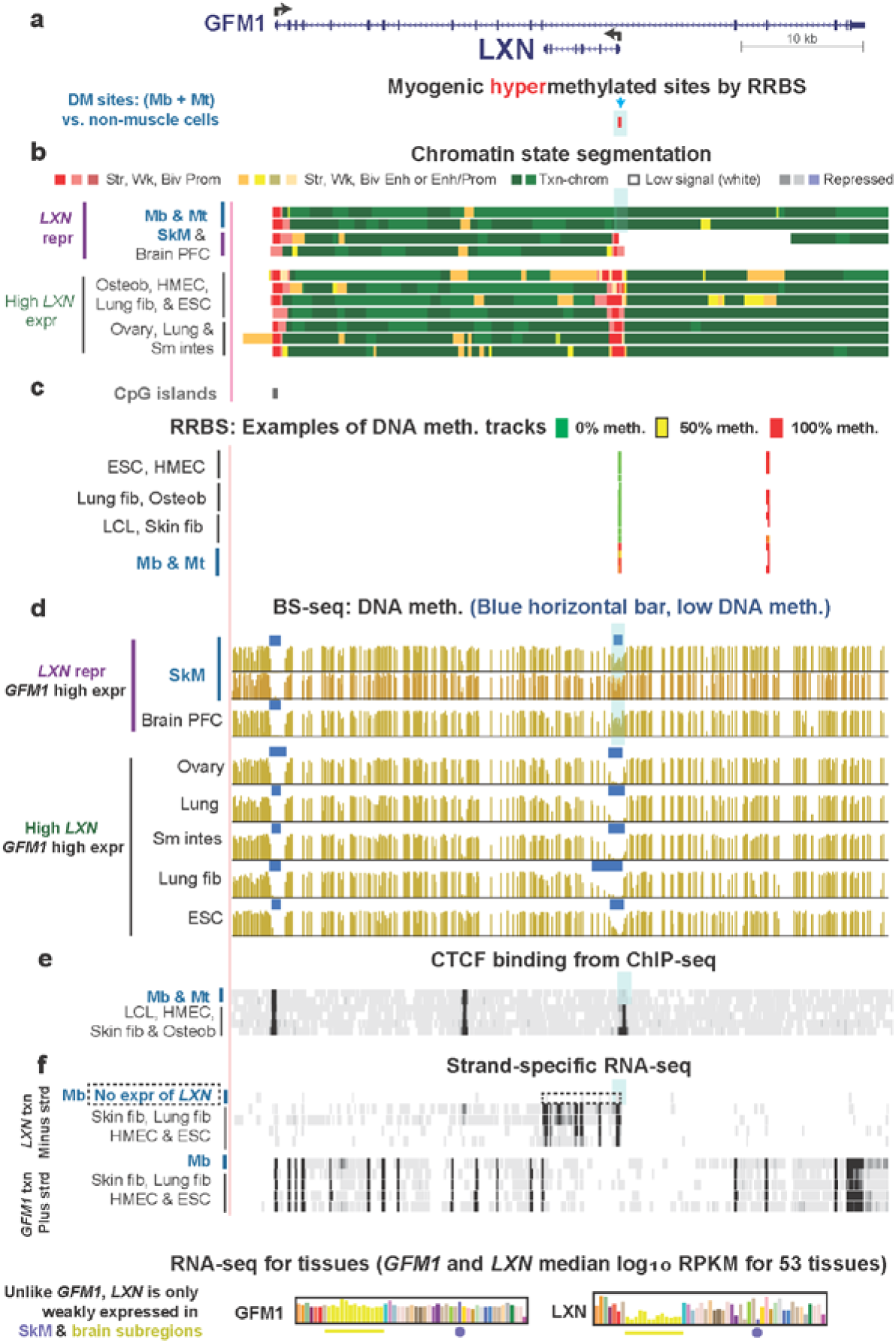
*LXN*, a tissue-specific gene within a constitutively expressed gene, displays promoter repression and DNA hypermethylation specifically in Mb but not repressive chromatin. **(a)** RefSeq gene structure for *LXN*, a carboxypeptidase-encoding gene, and *GFM1*, a gene encoding a translation elongation factor (chr3:158,358,796-158,412,265). Coordinates for gene figures are in hg19 from the UCSC Genome Browser ^34^ and all tracks other than GTex RNA-seq bar graphs are aligned. Also shown are statistically significant myogenic hypermethylated sites as determined by RRBS comparisons of the set of nine Mb and Mt samples vs. 16 types of non-muscle cell cultures ^28^; at this resolution individual CpGs cannot be seen. **(b)** 18-State chromatin state segmentation from RoadMap database ^23, 34^ with the indicated color code; Prom, promoter; Enh, enhancer; Enh/Prom, both active promoter-type and enhancer-type histone modifications; Repressed, enriched in H3K27me3 (weak, light gray; strong, dark gray) or H3K9me3 (violet). **(c)** CpG islands and examples of RRBS data tracks for DNA methylation levels at individual CpGs with a key for the semi-continuous color code; only some of the RRBS samples (and biological or technical duplicates) are shown. ^28^ **(d**) Bisulfite-seq profiles with blue bars indicating regions with significantly lower methylation compared to the rest of the given genome ^23, 33^. The two SkM samples are both from psoas muscle and are biological duplicates; the second one does not have scoring of its low-methylated regions with blue bars. **(e)** CTCF binding from chromatin immunoprecipitation/next-gen sequencing (ChIP-seq) indicating the Mb-and Mt-specific loss of CTCF binding from the DMR/promoter region. **(f)** Strand-specific RNA-seq shows no expression of *LXN* specifically in Mb while there is strong expression in the tested non-myogenic cell cultures. GTex RNA-seq data from hundreds of biological duplicates for each tissue type demonstrates only very low levels of *LXN* expression in SkM and in different subsections of brain and that *GFM1* is constitutively expressed. Fib, fibroblasts; osteob, osteoblasts; sm intes, small intestine. Blue arrow in (a), CCGG site tested in the Epimark assay to quantify 5hmC and 5mC levels; blue highlighting, the region of myogenic DNA hypermethylation.

**Figure 5.**
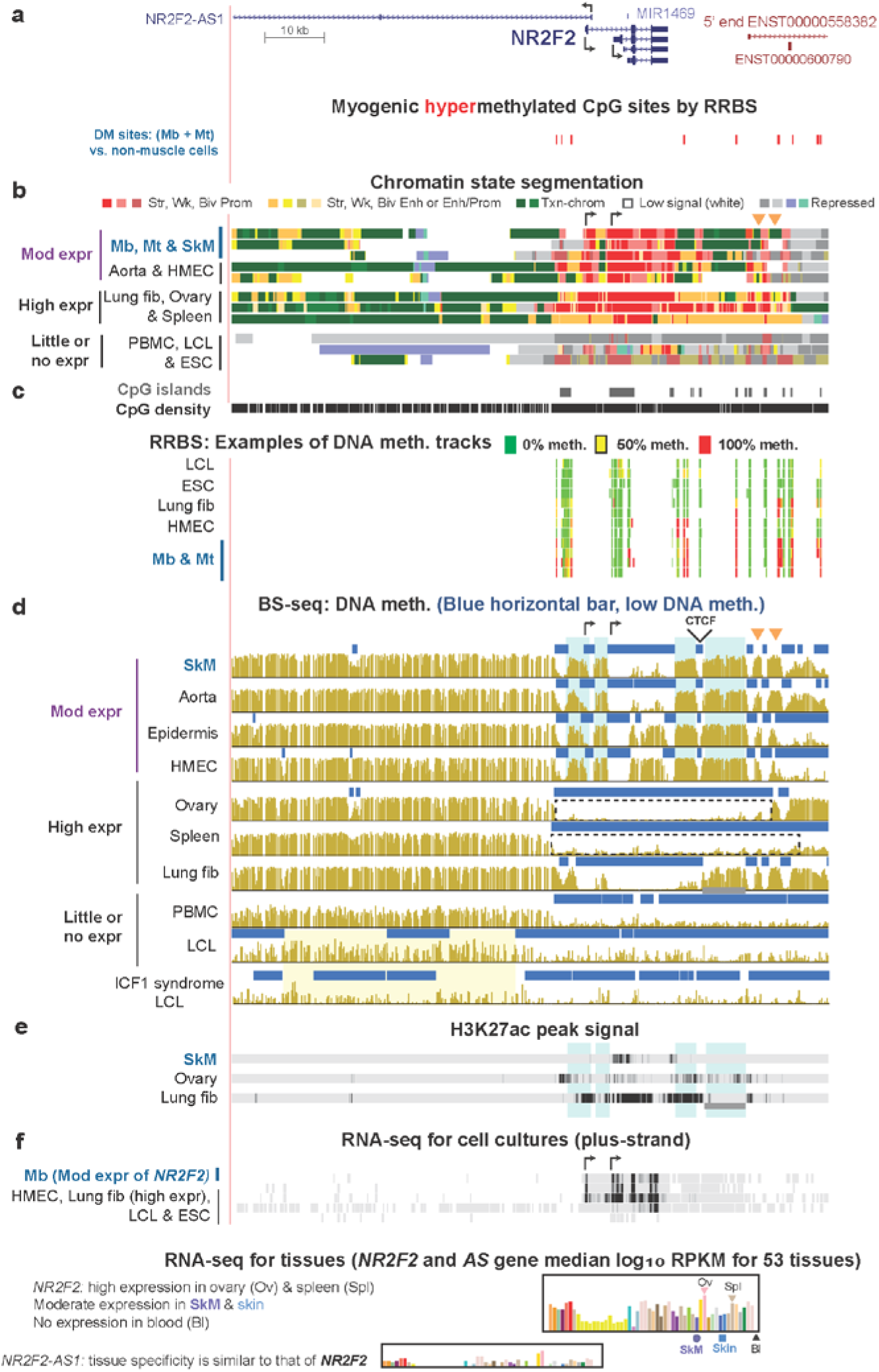
*NR2F2*, which encodes a master TF regulator, is down-modulated in Mb, SkM, aorta, epidermis and mammary epithelial cells and displays hypermethylated DMRs in those samples. **(a)** RefSeq structure for *NR2F2* and Mb-hypermethylated sites (chr15:96,808,300-96,911,119). **(b)** Chromatin state segmentation. Samples are divided into those with moderate, high, and little or no expression groups. The locations of two of the alternative TSS for *NR2F2* are indicated by broken arrows and the SkM-hypermethylated DMRs that correspond to gaps in promoter chromatin are shown by orange triangles. **(c)** Examples of RRBS tracks for cell cultures. **(d)** Bisulfite-seq profiles as in Figure 3 with the locations of two of the alternative TSS for *NR2F2*, a constitutive CTCF binding site, and two of the hypermethylated DMRs indicated above the SkM track. A second control LCL bisulfite-seq profile was available ^34, 43^ which gave similar results to the one shown. Dotted boxes, two of the super-enhancer regions that show very low levels of DNA methylation. **(e)** Enrichment in H3K27ac as in Figure 4x; gray horizontal line for lung fibroblasts in (d) and (f) shows the region of high DNA methylation and loss of H3K27ac that disrupts the super-enhancer in these cells. **(f)** RNA-seq.

**Figure 4.**
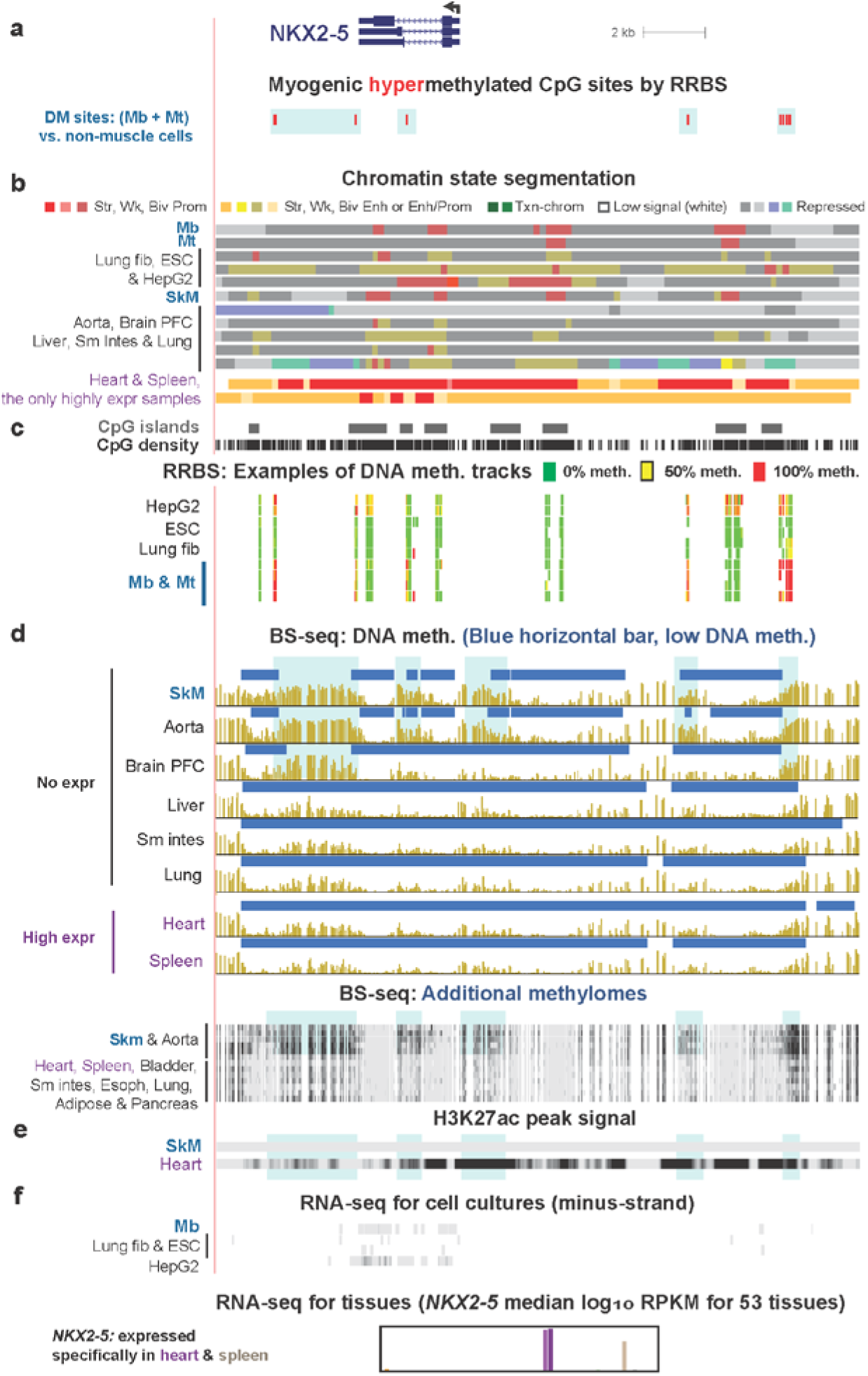
Cardiac TF-encoding *NKX2-5* is repressed in many samples without DNA methylation but exhibits repression accompanied by DNA hypermethylation in Mb, SkM, and aorta at a cryptic super-enhancer. **(a)** RefSeq structure for *NKX2-5* and Mb-hypermethylated sites as in Figure 3 (chr5:172,654,786-172,675,423). **(b)** Chromatin state segmentation as in Figure 1 with the addition that aqua green denotes enrichment in repressive H3K9me3 with low levels of H3K36me3. Heart and spleen, the only two high-expressing samples have extended enhancer-and promoter-chromatin regions that signify a super-enhancer extending from upstream to downstream of the gene. **(c)** Examples of RRBS tracks for normal cell cultures with the addition of the HepG2 liver cancer cell line. **(d)** Bisulfite-seq profiles as in Figure 3 with some additional bisulfite-seq tracks for other samples shown in the dense configuration to indicate the consistency of the SkM-and aorta-specific hypermethylation among replicates. Samples grouped as those with no expression and those with high expression of *NKX2-5*. **(e)** Enrichment in H3K27ac from peak calling using MACSv2 with a p-value threshold of 0.01 ^34^ to illustrate that several of the hypermethylated SkM DMRs overlap especially strong enhancer regions as shown by their overlap with strong H3K27ac signals. **(f)** RNA-seq as in Figure 1. HepG2, which like Mb had a very low level of expression, also shares the far upstream and gene-downstream hypermethylation with Mb. Esoph, esophagus; CpG density, plot of CpGs.

**Figure 6.**
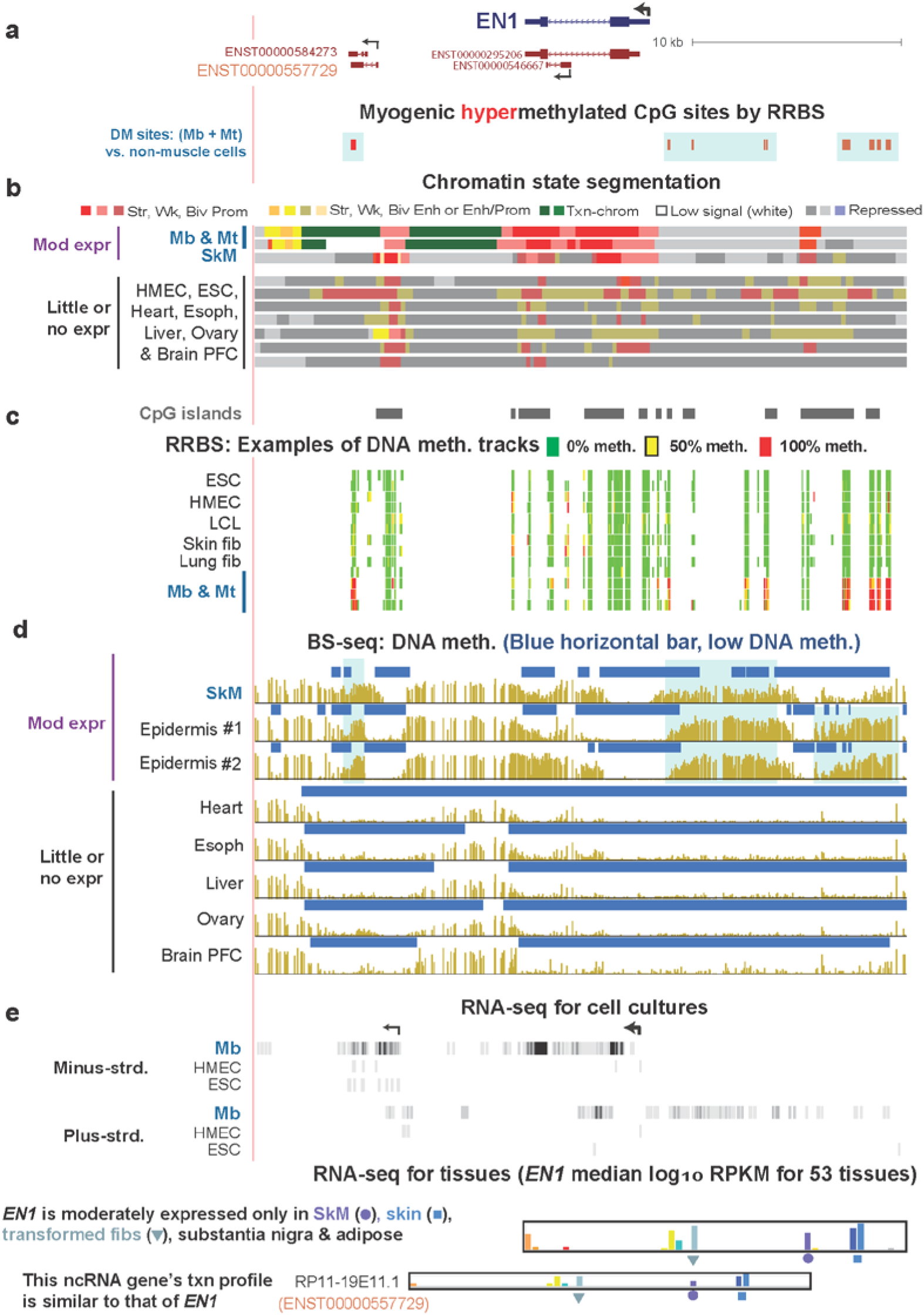
The homeobox gene *EN1* is preferentially expressed in Mb, SkM, and epidermis and has TSS-upstream and gene-downstream hypermethylation in those samples. **(a)** RefSeq structure for *NKX2-5* and Mb-hypermethylated sites (chr2:119,587,322-119,618,802). **(b)** Chromatin state segmentation. **(c)** Examples of RRBS tracks for cell cultures. **(d)** Bisulfite-seq profiles showing gene-upstream, gene-downstream, as well as 3′ intragenic hypermethylated DMRs in SkM and epidermis. **(e)** RNA-seq. The TSS for *EN1* and one of the downstream ncRNA genes are indicated by broken arrows.

Surprisingly, the most frequent tissue developmental association among the 95 Mb-hypermeth genes was with neural development. We found that 32 of these genes are implicated in embryonic development of the neural system, and only 21 are linked to SkM development (Supplementary Tables S1a-S4a). The only other frequent developmental association of gene function with Mb hypermethylation was with cardiac development (15 genes). With respect to sharing of T/C-specific hypermethylated DMRs, the strongest overlap between two sample types was for SkM and Mb with 74 of the 95 genes displaying SkM as well as Mb hypermethylation and usually with similar transcription status for the associated gene (Supplementary Tables S1b-S4b). There was also considerable overrepresentation of several non-myogenic sample types sharing DNA hypermethylation with Mb, namely, osteoblasts, aorta, human mammary epithelial cells (HMEC), skin fibroblasts and brain prefrontal cortex (PFC) for 30, 29, 22, 22 and 21 genes, respectively (Supplementary Tables S1b-S4b). In comparison, heart, ovary, liver, adrenal gland, skin, B-cell lymphoblastoid cell lines (LCLs), and small intestine samples exhibited hypermethylation at the Mb-hypermethylated DMRs at only 14, 7, 7, 6, 5, 4 and 2 genes, respectively. Twenty genes displayed DNA hypermethylation only in the SkM lineage.

### Analysis of hydroxymethylation at some Mb-hypermethylated sites

DNA methylation profiled by RRBS or bisulfite-seq cannot distinguish between 5hmC and 5mC. Therefore, we used an enzyme-based assay (Epimark) to quantify 5hmC and 5mC at specific sites ^36^. We assayed biological replicates of SkM, heart, cerebellum, leukocytes, spleen, lung, kidney, placenta, LCLs, non-transformed cell strains of Mb, skin fibroblasts and human umbilical vein endothelial cells (HUVEC) at three CCGG sites that were significantly hypermethylated. These sites are located 0.1 kb downstream of the TSS in *LXN* (TSS + 0.1 kb), 4.2 kb upstream of the TSS of *EBF3* (TSS – 4.2kb) and in *SIM1* (TSS + 0.1kb; blue arrows in Figure 3a & Supplementary Figure S7a and e). *LXN* is repressed and *EBF3* is preferentially expressed in Mb and SkM. *SIM1* is moderately expressed in Mb and mostly repressed in SkM. SkM had appreciable 5hmC only at the *EBF3* and *SIM1* sites (26 and 11% of C as 5hmC, respectively), which had more than twice as much 5mC as 5hmC (Supplementary Table S5a). As we found at other Mb DM sites that we examined in Epimark assays in earlier studies ^36^, there were generally only negligible levels of 5hmC in cell cultures, leukocytes, spleen, lung, placenta and sperm.

A hydroxymethylome profile by TAB-seq ^34, 37^ for the examined tissues or postnatal cell cultures is available only for brain PFC. While a comparison of bisulfite-seq (detecting 5mC and 5hmC) and Tab-seq (detecting just 5hmC) on the same DNA sample is not strictly quantitative, it can indicate whether there is much or little 5hmC relative to 5mC in a given region for a particular sample. Such a comparison for genes that were methylated in brain as well as in Mb at the Mb-hypermethylated DMRs revealed that 16 genes had much more 5mC than 5hmC over the brain PFC DMR (*SIX3* and *SIX2*, Supplementary Figure S2e and Table S5b). Nine genes had considerable levels of both 5hmC and 5mC at the DMRs (*ZIC4*, Supplementary Figure S3e and f and Table S5b).

### Most genes displaying DNA hypermethylation and repression in myogenic cells were also silenced in other cell types but by DNA methylation-independent mechanisms

Of the 31 Mb-hypermeth/repr genes, 26 had their DMR within 2 kb upstream or downstream of the TSS, and 18 of these promoter-DMRs were very densely methylated as indicated by their being located in a CGI. However, only five of the 31 Mb-hypermeth/repr genes (*LXN*, *MARVELD2, GALNT6, ESRP2* and an unannotated lncRNA gene near *SMIM21;* Supplementary Table S1a) displayed DMR hypermethylation in most or all the cell cultures or tissues in which the DMR-associated gene was repressed. Repression without DNA methylation in non-myogenic samples was associated with PcG-chromatin for 23 of the genes (Supplementary Table S1a). *LXN*, one of the five genes whose repression was strictly associated with promoter DNA methylation, is of particular interest because the tight linkage of its repression to promoter hypermethylation is probably related to its unusual location. This small gene, which encodes an inflammation-associated carboxypeptidase inhibitor, is embedded in intron 13 of a *GFM1*, a large constitutively expressed gene (Figure 3). In Mb, the *LXN* promoter-region DMR is embedded in txn-chromatin (Figure 3b). A likely explanation for the use of DNA methylation and not repressive chromatin at the *LXN* DMR to silence this gene in Mb is that if there were repressive PcG-chromatin at the *LXN* promoter, which is intragenic to *GFM1*, it probably would have interfered with *GFM1* expression.

Some of the Mb-hypermeth/repr genes that were repressed in many non-myogenic cell cultures or tissues without DNA hypermethylation had gene neighbors that were preferentially expressed in myogenic cells, e.g., *SIX3* and *SIX2* (Supplementary Figure S2). *SIX3* is silent in almost all studied myogenic and non-myogenic samples but hypermethylated at DMRs upstream and downstream of the gene and within its single intron in Mb, Mt, SkM and aorta. These samples specifically express *SIX2*, the closest protein-encoding gene to *SIX3* albeit 59 kb downstream. Brain pre-frontal cortex DNA shares several of these hypermethylated *SIX3* DMRs although it does not express *SIX2*. Hypermethylation around *SIX3* in brain PFC might be related to this gene’s selective expression in brain basal ganglia (Supplementary Figure S2a). DNA methylation could be needed in addition to PcG-chromatin silencing and differential TF activity to efficiently repress this gene in a non-expressing part of the brain. Similar examples of neighboring pairs of a Mb-hypermeth/repr gene and a Mb preferentially expressed gene are *SIX6* and *SIX1*; *PNMA8B* and *PNMA8A; ZIC4* and *ZIC4; HSD17B14* and *PLEKHA4* (Supplementary Tables S1-S3; Supplementary Figure S3). The hypermethylation of the repressed gene of these gene-pairs might be necessary because of the extremely close distance between the two genes (0.5 kb, *HSD17B14* and *PLEKHA4*) or the proximity to the repressed gene of the expressed gene’s enh-chromatin (Supplementary Figure S2c) ^38^ or its own promoter, which sometimes can act as an enhancer. Interestingly, *SIX2* and *ZIC1* genes themselves have a positive association of transcription with DNA methylation while their neighbors, *SIX3* or *ZIC4*, have a negative association (Supplementary Figures S2b and S3a).

Two of the Mb-hypermeth/repr genes *NKX2-5* and *IRX4* were expressed specifically in heart plus one or a few other tested T/C types (spleen for *NKX2-5* and esophageal mucosa, skin, and HMEC for *IRX4*; Figure 4 and Supplementary Table S1). Hypermethylation of the *NKX2-5* DMRs was seen in aorta and the HepG2 liver cancer cell line as well as SkM and Mb. In heart and spleen, where these DMRs were not hypermethylated, they were embedded in a 20 – 23 kb region of enh-chromatin (super-enhancer or stretch enhancer ^18, 39^) that was interspersed with prom-chromatin spanning from upstream to downstream of this small gene (2 kb; Figure 4b).

### Myogenic DNA hypermethylation associated with down-modulated expression was often located in cryptic enhancers

Twenty-two of the 31 Mb-hypermeth/downmod genes had DMRs that overlapped unmethylated or weakly methylated strong enh-chromatin in non-myogenic, highly expressing samples (Figure 2d; Supplementary Table S2a and b). Importantly, only one or a few diverse, non-myogenic samples (e.g., spleen, lung, brain, esophagus, adipose, ovary, skin fibroblasts or osteoblasts) exhibited enh-chromatin at the Mb-hypermeth DMRs so that without examining many sample types these enhancers could easily be missed. ChIP-seq TF-binding databases were mostly uninformative about the enh-chromatin in this study because Mb were excluded from most available genome-wide profiles.

*NR2F2 (COUP-TFII)*, is a Mb-hypermeth/downmod gene which is particularly influential during development and has complicated T/C-specific epigenetics. It encodes a TF with key roles in many types of development, including myogenesis, cardiovascular development & neurogenesis, as well as in metabolic homeostasis and disease (especially cancer progression and metastasis) ^40^. Like eight other genes that we analyzed (Supplementary Tables S1a-3a), it regulates the epithelial-to-mesenchymal transition ^41^. Mb, SkM, skin and HMEC display intermediate levels of expression and had hypermethylated DMRs that are upstream and downstream of the gene (Figure 5d). In highly expressing, non-myogenic samples, *NR2F2* and nine other Mb-hypermeth/downmod genes had a long hypomethylated super-enhancer in one or more high-expressing non-myogenic samples that overlaid the Mb DMRs (Supplementary Figures S6 and S8 and Table S2a). Ovary, spleen and lung fibroblasts highly express *NR2F2* and displayed a long super-enhancer spanning the gene and beyond (Figure 5b). In contrast, peripheral blood mononuclear cells (PBMC), an LCL, and ESC exhibited little or no expression of this gene; however, they too had only little DNA methylation at the DMR cluster (Figure 5c and d). The methylation profile in and around DMRs in repressed samples for Mb-hypermeth/downmod genes often displayed more scatter and less well defined borders than did DMRs in high-expressing samples (Figure 5d and Supplementary Figure S8d). Although *NR2F2* did not show DMR/alternative promoter correlations, it did illustrate the relationship between DNA hypermethylation and the detailed shaping of prom-and enh-chromatin regions. Hypermethylated DMRs downstream of *NR2F2* in many of the expressing samples (Figure 5d, orange triangles) interrupt a region of prom-chromatin possibly to silence lncRNA gene expression (Figure 5a and b). Similar DNA hypermethylation-associated shaping of promoter or enhancer regions was seen downstream of the *NR2F2* super-enhancer in lung fibroblasts (Figure 5d and e, gray horizontal bars) and upstream of the TSS of *EBF3* in Mb, SkM, and heart (Supplemental Figure S7a-c, pink highlighting). In addition, upstream of *NR2F2* was a disease-related hypomethylated DMR associated with the ICF1 syndrome (immunodeficiency, centromeric region instability, facial anomaly), a rare recessive disease that results from loss of most DNMT3B activity. In our previous transcriptome analysis of many ICF and control LCLs, one of the most significantly upregulated genes was *NR2F2* ^42^. Upstream of *NR2F2* and overlapping *NR2F2 AS1* (Figure 5d, yellow highlighting), there was a long region of DNA hypomethylation in a *DNMT3B*-mutant ICF1 LCL relative to two controls ^23, 34, 43^ suggesting that this hypomethylated DMR may be responsible for the qRT-PCR-confirmed upregulation of *NR2F2*^42^ in ICF vs. control LCLs.

### Intergenic or intragenic myogenic DNA hypermethylation associated with genes preferentially expressed in myogenic cells

Of the 20 Mb-hypermeth/pref-expr genes, 12 had Mb hypermethylation upstream or downstream of the gene indicating DNA methylation that was not just the previously described gene-body DNA methylation often associated with transcription elongation (Supplementary Table S3) ^23^. *EN1* illustrates a gene with preferential transcription in Mb as well as in SkM and epidermis that displayed such gene-upstream and downstream hypermethylation in these samples (Figure 6a, c and d). It specifies a homeobox TF found in the dermomyotome during embryogenesis. Mb, SkM, and skin had hypermethylated DMRs 0.9 kb upstream and 14 kb downstream of the gene immediately distal to the promoter chromatin in Mb, Mt and SkM (Figure 6a-d). This suggests that border-like DNA hypermethylation upstream of the gene suppresses the spread of the promoter-adjacent repressive chromatin into the promoter ^5, 44^. In addition, both upstream and downstream of the promoter (Figure 6e, minus-and plus-strands and RNA-seq for tissues), Mb hypermethylation may also aid transcription of long-lived AS or sense ncRNAs that might facilitate *EN1* transcription.

*SIX2* is another example of Mb-hypermeth/pref-expr gene encoding homeobox TF. It is very highly expressed in Mb and moderately expressed specifically in SkM and aorta. The hypermethylated DMR in these samples starts at the 3’ end of the gene and overlays txn-and weak prom-chromatin in Mb and Mt (Supplementary Figure S2). This Mb/SkM/aorta DNA hypermethylation might facilitate *SIX2* expression by counteracting the spread of gene-downstream, repressive H3K9me3-enriched and PcG-chromatin into the prom-chromatin that covers almost the entire gene. Similarly, *SIM2* and *TBX18*, Mb-hypermeth/pref-expr genes which also encode developmental TFs, displayed Mb DNA hypermethylation immediately upstream of their promoters adjacent to repressive PcG-chromatin (Supplementary Table S3).

### Intergenic or intragenic myogenic DNA hypermethylation associated with repressed alternative or cryptic promoters

*ZIC1*, which encodes a neurogenic and myogenic TF ^45, 46^, like some other genes with Mb-hypermethylated DMRs, displayed associations between DNA hypermethylation and the choice of alternative promoters. DNA hypermethylation at several regions upstream and downstream of *ZIC1* in Mb, SkM, osteoblasts and skin fibroblasts was associated with use of a previously undescribed alternative promoter for *ZIC1* within intron 3 of the adjacent and oppositely oriented *ZIC4* gene (Supplementary Figure S3a and b, large arrow). *LAD1*, another Mb-hypermeth gene associated with alternative promoter use, encodes an epithelial membrane protein and has a hypermethylated and repressed canonical promoter in Mb. In Mb an intragenic cryptic promoter gives rise to a highly 5’-truncated RNA (Supplementary Figure S5d, blue box). Mb DNA hypermethylation at the canonical *LAD1* promoter is probably related to *LAD1* having 5’ and 3’ gene neighbors (*TNNT2* and *TNNI1*) that are preferentially expressed in Mb and Mt and to the gene body of *LAD1* overlapping a myogenic super-enhancer that spans *LAD1* ^47^. The intragenic *LAD1* lncRNA might contribute to this myogenic superenhancer activity for *TNNT2* and *TNNI1*. *TBX1* is also predominantly expressed from a cryptic intragenic promoter. Its DNA methylation in the 1-kb upstream region could not be ascertained in our previous RRBS study due to the fact that RRBS covers only a small, but usually informative, subset of CpG sites ^20^. From recently available bisulfite-seq profiles of SkM samples ^23^ it can be seen that there is dense SkM-lineage-specific methylation in this canonical promoter region (Supplementary Table S3a). Both Mb and SkM strongly and specifically express this gene but have active promoter chromatin only in the middle of the gene body (Supplementary Table S3a).

The *DBX1* homeobox gene is a Mb-hypermeth/repr gene with a 3’ DMR that overlaps a cryptic promoter for an ncRNA expressed specifically in ESC (Supplementary Figure S4e, blue box). The DNA hypermethylation targeted to myogenic cells may be necessary to silence this ncRNA promoter in myogenic cells irrespective of *DBX1* transcription. In contrast, Mb hypermethylation and repression of intragenic cryptic promoters of *JSRP1* (Supplementary Figure 10)*, STAC3, CDH15, PITX3*, and *RYR1* were positively associated with expression of these genes in myogenic cells. Their DMRs were embedded in weak or bivalent promoter chromatin at the cryptic promoter in non-myogenic samples in which RNA-seq and 5’ cap analysis of gene expression (CAGE) profiles indicated that the unmethylated DMRs were capable of acting as promoters in vivo (Supplementary Figure S10c and d, dotted boxes, and Supplementary Table S2a-4a). The cryptic promoters when unmethylated were associated with bidirectional transcripts (*CDH15* and *PITX3* ^36^), AS RNA (*RYR1*), or sense transcripts (*JSRP1* and *STAC3*). For these five genes, Mb hypermethylation is probably silencing an intragenic ncRNA promoter that would otherwise interfere with mRNA generation in Mb.

### Inhibition of binding to CTCF sites at Mb-hypermethylated DMRs

Myogenic hypermethylation at DMRs was associated with decreased binding of the CCCTC-binding factor (CTCF) near 13 of the 95 examined genes (Supplementary Tables S1a-S4a), as determined from CTCF ChIP-seq profiles, which are available for 12 of the RRBS-analyzed cell cultures. CTCF can act as a DNA methylation-sensitive TF, mediate insulation, modulate alternative splicing, and cause changes in higher-order chromatin structure that affect transcription initiation^48^ ^49^. An example of a Mb-hypermeth gene with promoter and CTCF site hypermethylation is the above-described *LXN* (Figure 3f). Mb and Mt were uniquely lacking in binding of CTCF to the 5’ end of *LXN* and their highly specific Mb-hypermeth DMR overlaps this site (Figure 3e). However, the predicted binding sequence at this site does not contain any CpG sequences, unlike many CTCF sites (CTCF ChIP-seq, ^34^). Therefore, methylation-blocked CTCF binding is probably contributing to *LXN* promoter shutdown in myogenic cells indirectly, e.g., by limiting promoter availability. A different type of correlation of CTCF binding and transcription was seen at the 3’ end of *LBX1-AS1*, which shares a bidirectional promoter region with *LBX1*. Loss of CTCF binding to this binding sequence, which contains two CpGs, was seen at a SkM-lineage hypermethylated DMR (Supplementary Figure S9a and f). This decreased binding of CTCF at the 3’ end of *LBX1-AS1* and other nearby changes in CTCF binding were positively associated with transcription of *LBX1*, suggesting that the DMR causes transcription-modulating alterations in chromatin conformation in this gene neighborhood. In *NR2F2*, constitutive binding of CTCF downstream of the gene in the middle of a Mb/SkM/epidermis/HMEC hypermethylated DMR was seen in all studied cell cultures. The region around the CTCF site constitutively lacks DNA methylation providing an unmethylated valley in the middle of T/C-specific hypermethylated DMR (Figure 5d).

## Discussion

A comparison of epigenetic and transcription profiles of 95 genes associated with myogenic hypermethylated DMRs in 38 types of samples revealed four major categories of DNA hypermethylation/transcription relationships. They are gene silencing (Mb-hypermeth/repr genes); transcription down-modulation rather than repression (Mb-hypermeth/downmod genes); preferential transcription (Mb-hypermeth/pref-expr genes), and regulation of the use of alternative or cryptic promoters (Figure 2d). Even though the 95 genes were chosen only based on discernible relationships between myogenic DNA hypermethylation and transcription (Figure 2d), 70 of them encode development-associated proteins, mostly developmental TFs. A genome-wide relationship of all myogenic hypermethylated CpGs to early development was also seen in the strong enrichment for the gene ontology terms “DNA binding” and “embryo development” and the frequent overlap of the global Mb-hypermeth sites with ESC-specific bivalent chromatin (Figure 2b).

Most of the 31 Mb-hypermeth/repr genes had their DMRs in the upstream promoter region or in the region immediately downstream of the TSS, a result consistent with previous findings about TSS-adjacent repressive DNA methylation^2, 50^. Some Mb-repr genes displayed both promoter-region and non-promoter hypermethylation in Mb suggesting that promoter hypermethylation did not suffice for silencing transcription in those cells. Surprisingly, only five of the Mb-hypermeth/repr genes displayed a tight association of DNA hypermethylation with repression (Figure 7b). The other 26 Mb-hypermeth/repressed genes are repressed in many non-myogenic samples without DNA hypermethylation but usually with repressive H3K27me3-enriched chromatin (PcG-chromatin) in cis (Figure 7a). Therefore, the Mb DNA hypermethylation was not simply the consequence of silencing these genes. Some of the observed DNA hypermethylation in postnatal Mb might persist as a kind of epigenetic memory just reflecting a history of DNA methylation involvement in transcription control during prenatal development ^22, 51^. However, for many early-development genes, our analysis suggests that this DNA hypermethylation is functional, which is consistent with recent studies of DNA hypermethylation^14, 24, 52^. Certain Mb-hypermeth/repr genes involved in specifying non-myogenic cell lineages may require DNA methylation for efficient gene repression in Mb and SkM because of a need to guard against even very low levels of their expression in the SkM lineage. For example, DNA hypermethylation throughout the *NKX2-5* vicinity in Mb, SkM, and aorta is linked to the absence of a heart-and-spleen super-enhancer although most tissues and cell cultures silence this cardiogenic TF-encoding gene without DNA hypermethylation. DNA hypermethylation of *NKX2-5* may have to be targeted to the SkM lineage because of the strong overlap of gene expression profiles from heart with those from SkM ^53^ and the partial sharing of cardiogenic and myogenic developmental TFs (cardiogenic and facial SkM myogenic TFs^54^). Indeed, inappropriate expression of *NKX2-5* in SkM of myotonic dystrophy type 1 patients or in transgenic mice SkM or stably transfected C2C12 Mb is linked to interference with normal SkM development ^55^. Furthermore, we propose that the previously reported *NKX2-5* intragenic, disease-linked gene-body DNA hypermethylation, which was coupled with lower *NKX2-5* mRNA levels in the hearts of some cardiac patients with tetralogy of Fallot,^56^ involves decreased super-enhancer formation.

**Figure 7.**
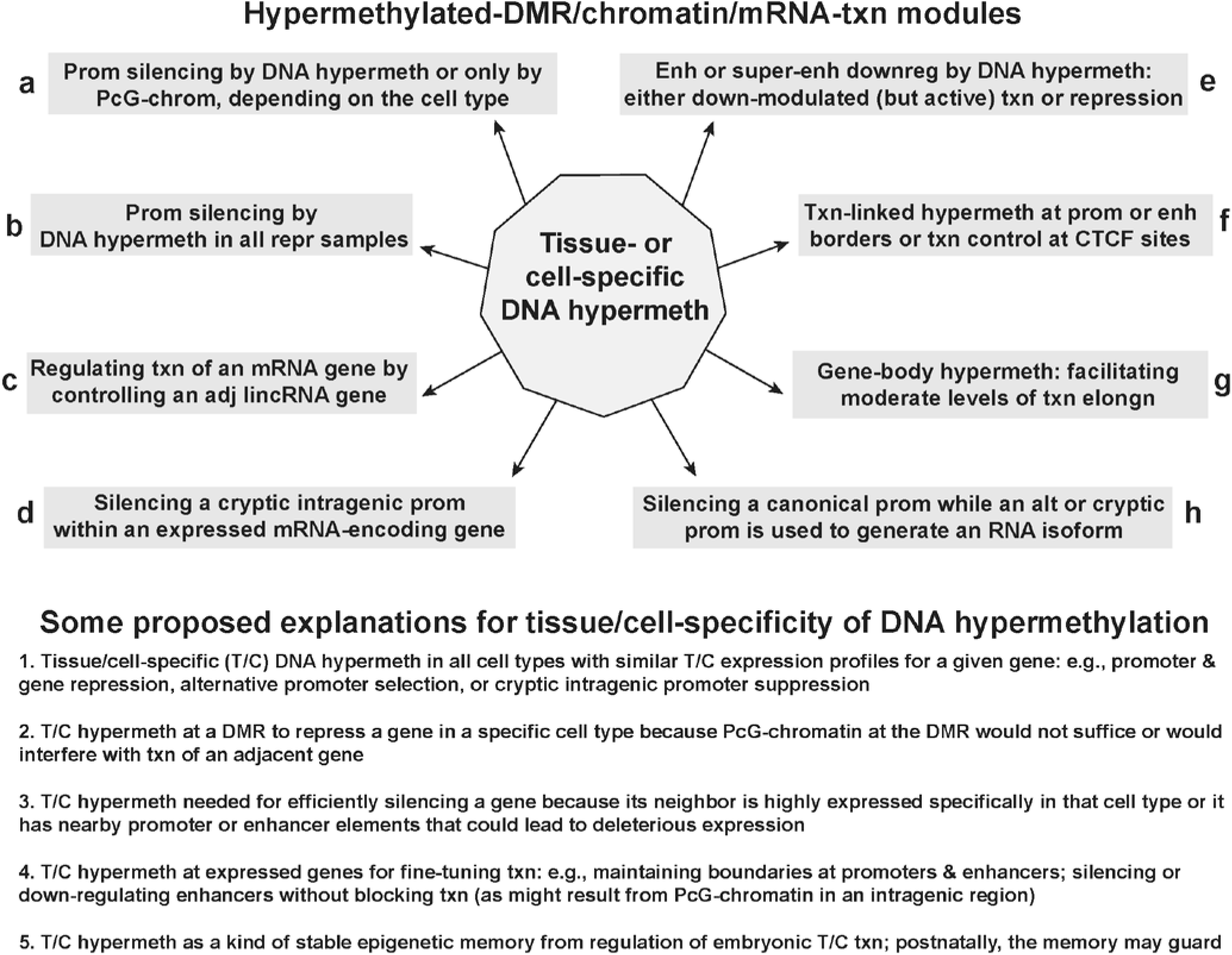
Models for relationships between hypermethylated DMRs, chromatin state, and transcription of the associated mRNA-encoding genes for the 95 highlighted Mb-hypermethylated genes. **(a)** through **(h)** summarize models for the interconnections between DNA hypermethylation, chromatin epigenetics, and transcription of the 95 protein-encoding genes, which were mostly development-associated genes, that were examined in detail in this study. **(g)** Gene-body hypermethylation may facilitate moderate levels of transcription by suppressing the formation of unstable ncRNA transcripts in contrast to the stable ncRNA transcripts referred to in **(d)**. This list of functions for DNA methylation reflects the associations that we saw among the 95 examined genes and is not meant to all-inclusive for DNA methylation-transcription relationships. For example, the association of alternative splicing to DNA methylation is not included although four of the 95 genes gave some evidence for a correlation between T/C-specific DNA hypermethylation and alternative use of splice isoforms (Supplementary Table S4a). In the bottom part of the figure, five of the proposed biological rationales for these differentially methylated regions (DMRs) are given. Hypermeth, hypermethylation; txn, transcription; enh, enhancer; prom, promoter; PcG-chrom, chromatin enriched in repressive H3K27me3; adj, adjacent; lincRNA, long intergenic noncoding RNA; downreg, down-regulation; T/C, tissue-or cell type-specific; elongn, elongation; “at CTCF sites” includes sites near but not within the consensus CTCF binding sequence.

For the 31 Mb-hypermeth/downmod genes, there were both positive and negative correlations of DNA hypermethylation with gene expression. These genes are moderately expressed in Mb and often also in several non-myogenic samples in association with DMR hypermethylation. However, DMR methylation was lacking in both non-myogenic samples in which the genes are more highly expressed and in samples in which they are not expressed. Most of these genes had DMRs that overlapped enhancers or super-enhancers that were active (enriched in H3K27ac and H3K4me1) specifically in one or a few non-myogenic lineages in which the genes were highly expressed. Therefore, DNA methylation for these genes appears to be linked to down-regulating enhancers in cis and thereby down-modulating transcription in cell cultures and tissues that display moderate expression of the gene (Figure 7e). The seemingly paradoxical lack of DMR methylation at cryptic enhancer/DMRs in non-expressing samples for Mb-hypermeth/downmod genes can be explained by repressive chromatin sufficing for epigenetic silencing of the enhancers in these genes when the promoter is inactive. It is noteworthy that cryptic super-enhancers in repressed genes often had more scattered partial methylation throughout the DMR and at its borders than did active enhancers (e.g., Figure 5d). Active enh-chromatin generally exhibits low DNA methylation for at least part of its length ^6, 39, 57, 58^, and this hypomethylation is implicated in enhancer formation ^52^. Similarly, we previously demonstrated that in vitro methylation targeted to only the three CpGs within the powerful 258-bp core enhancer of the 40-kb SkM-lineage-specific *MYOD1* super-enhancer sufficed to decrease enhancer activity by almost 90% in reporter gene assays in Mb ^58^. Moreover, these three CpGs are hypomethylated in SkM where the enhancer is active. Our genome-wide Mb-hypermethylation profiling showed an inverse association of gene-body DNA methylation and the strength of enhancer activity as discerned from histone modifications (Figure 1d), which suggests that DNA methylation can fine-tune enhancer activity as well as silence it. Despite the importance of enhancer DNA hypomethylation, recent evidence indicates that DNA methylation in certain enhancer subregions actually promotes enhancer activity ^57, 59^. However, we found that some gene-spanning super-enhancers in highly expressing samples had only small subregions with DNA methylation (*NR2F2*, Figure 5d and *TBX3*, Supplementary Figure S8d, dashed boxes) suggesting that for some super-enhancers either small subregions of DNA methylation suffice or some enhancers do not require methylated subregions.

The epigenetics of *NR2F2*, a Mb-hypermeth/downmod gene that codes for a master regulatory TF for organogenesis and physiology, is of particular interest. The importance of the *NR2F2-*upstream hypermeth-DMR seen in Mb, SkM, aorta, HMEC and epidermis to fine-tuning *NR2F2* transcription (Figure 7c) is suggested by a comparison of ICF1 syndrome (DNMT3B-deficiency) and control LCLs. *NR2F2* is upregulated several fold in ICF LCLs vs. comparable control LCLs ^42^ and, in this study, was shown to have a long, mostly unmethylated ICF-associated DMR that overlapped the upstream lncRNA gene (Figure 5d). Therefore, abnormal DNA hypomethylation-linked upregulation of *NR2F2* in lymphocytes might contribute to ICF immunodeficiency. *NR2F2* dysregulation has also been associated with exacerbating muscular dystrophy symptoms ^60^ and with carcinogenesis and metastasis ^41^.

One of the 20 Mb-hypermeth/pref-expr genes, *CDH15*, is very highly expressed in Mb and not in any non-myogenic cell cultures and had high levels of DNA methylation throughout most of the gene-body (exons 2-14) in Mb (Supplementary Table S3a) ^36^. This finding indicates that the intragenic DNA hypomethylation that we observed in other very highly expressed genes, like *NR2F2* in ovary and spleen, is not simply a default state due to dense packing of the transcription elongation machinery making the intragenic DNA inaccessible to DNMTs ^61^. We previously showed that *CDH15* gene-body methylation in Mb includes T/C-specific hypermethylation at a cryptic promoter that functions as a strong bidirectional promoter in C2C12 Mb but only when it is experimentally demethylated ^36^. *PITX3* ^36^ and *RYR1* are two other Mb-hypermeth/pref-expr genes for which intragenic hypermethylation appears to be linked to loss of an inhibitory gene-body promoter. Moreover, 35 of the 41 genes that were expressed in Mb and contained intragenic Mb-hypermeth DMRs exhibited overlap of these DMRs with such cryptic promoters or with alternative promoters (Figure 7h) or cryptic enhancers (Figure 7e). This finding is consistent with reports that transcription-associated intragenic DNA methylation (Figure 7g) is often important for suppression of interfering transcription initiation-regulatory sequences located in gene bodies ^1, 3, 13, 14^. The finding of expressed genes with DNA hypomethylation and extended enhancer or promoter chromatin throughout most of the gene body (e.g., *NR2F2* in ovary and *NKX2-5* and *EN1* in heart, Figures 4–6) raises the question of whether such chromatin minimizes the deleterious activation of cryptic intragenic promoters. Moreover, hypermethylation in the single intron of *NKX2-5* and x*SIX3* in the SkM lineage, where these genes are repressed, exemplifies gene-body hypermethylation associated with repression and suppression of a cryptic enhancer rather than with expression (Figure 7g).

Mb-hypermeth DMRs upstream of the core promoter region were observed for nine Mb-hypermeth/pref-expr genes (including *EN1*, *IRX3, TBX18* and *EBF3*). Although 5hmC upstream of promoters can be positively associated with transcription ^44^, we found negligible 5hmC in Mb at a tested DM site upstream of the TSS of the Mb-hypermeth/pref-expr *EBF3* gene as well as at CCGG sites associated with nine other genes examined in this study (Supplementary Tables S3a, S4a and S5a). We propose that promoter-upstream 5mC-rich DMRs associated with these nine genes help form a border ^4, 23, 44^ that limits the outward spread of prom-chromatin and the inward spread of repressive PcG-chromatin (Figure 7f).

Finding a biological rationale for the T/C specificity of DNA hypermethylation often requires considering the contexts of the gene’s neighborhood (neighboring genes & enhancer or promoter elements), the gene’s developmental history, its possible role in an inducible process like SkM repair, and its T/C-specific gene expression patterns among different cell lineages. *LAD1/TNNT2* /*TNNI1* in Mb and Mt and *SIX3/SIX2* in Mb, Mt, SkM, osteoblasts and aorta were hypermethylated and repressed selectively in cell/tissue types in which a neighboring gene is highly and specifically expressed. Other key developmental genes need fine-tuning of transcription, and not simply silencing, depending on the tissue, stage of development, or physiological needs. *TBX1* is a major player in the formation of many facial and neck skeletal muscles ^54^. It may influence SkM type postnatally ^62^ and, like *NKX2-5*, encodes a TF required for formation of the secondary heart field ^63^. *TBX1* haploinsufficiency is linked to the heart and skeletal muscle defects of the DiGeorge and velocardialfacial syndromes ^63^ indicating that the levels of its protein product must be tightly controlled. Other genes in this study (e.g., *TBX3, TBX4, TBX18, SIM1, ZIC1, NR2F2* and *PITX1*) require precise regulation of expression as seen in their linkage to haploinsufficiency-caused disease in humans, to transgenic mouse models of human disease, or in their response to environmentally-associated disease ^64^. *TBX1* is also implicated in interconversions of adipocyte subtypes ^65^. It has a densely methylated inactive canonical promoter and a novel unmethylated active promoter in the gene body in SkM and Mb. ^36^ Alterations in DNA methylation in and around this gene might modulate gene expression during embryonic development of facial/neck SkM vs. postnatal limb and trunk SkM, adipocyte interconversions, and physiological changes in SkM physiology. Examples of genes with changes in intragenic DNA methylation during SKM development (Mb vs. SkM tissue) that correlate with very different levels of gene expression are provided by *CORO6*, a cytoskeletal actin filament-encoding gene, and *STAC3*, a gene encoding a component of the excitation-contraction machinery of SkM (Supplementary Tables S2a and b).

There is remarkable diversity of the nonmyogenic differentiation pathways associated with many of the examined Mb-hypermeth genes. This diversity likely contributes to the need for hypermethylated DMRs to fine-tune expression for different developmental fates. Twelve of the analyzed 95 genes are involved in both embryonic myogenesis and neurogenesis (*PAX3, PAX7, SIM1, SIM2, ZIC1, TWIST1, EBF3, LBX1, NRXN2, EN1, LHX2*, and *KCNQ4*; Supplementary Tables 2a-4a). Five Mb-hypermeth genes are implicated in directing both myogenesis and adipogenesis (*TBX1, ZIC1, EN1, EBF3*, and *TCF21*), or in Mb transdifferentiation to adipocytes (*PRDM16* ^66^). Such genes may be more likely than most to require T/C-specific DMRs to differentially regulate their expression depending on temporal and spatial factors.

Many of the 95 studied genes genetically interact with one another during embryogenesis suggesting a kind of developmental co-methylation for fine-tuning their expression ^67^, e.g., *Tbx1*, which is genetically upstream of *Tcf21* and *Lhx2* in facial muscle development in the mouse ^54,68^. In addition, *PAX3* is implicated in the up-or down-regulation of many of the studied genes during SkM formation from somites or limb bud: *PAX7, SIM1, ZIC1, TWIST1, DBX1, TBX3, DMRT2, MEIS1*, and *GBX2* (Supplementary Tables S2a and 3a) ^69,70^. *PAX3* and *PAX7* not only regulate prenatal myogenesis but also postnatal muscle satellite cell renewal, induction of satellite cells to form Mb, fusion of Mb with myofibers during local SkM repair, and determination of SkM subtype ^26,71,72^. Therefore, these genes may need to be poised epigenetically for induction (Figure 7, bottom). In accord with their multiple roles in development, *PAX3, PAX7, TBX1*, and *NR2F2* have been shown to require different concentrations of their encoded TFs at different times and in different lineages in development ^63,73–75^, a need that can be fulfilled in part for these and many of other key developmental genes that we studied by differential DNA methylation. The obverse side of differentiation is cancer formation. Indeed, cancer-linked up-or down-regulated expression of many of the studied Mb-hypermeth differentiation-determining genes (e.g., *PAX3*, *NR2F2*, and *TWIST*) is implicated in carcinogenesis ^41,76^.

### Conclusions

There are many ways that differential DNA methylation can help regulate gene expression during and after development. We have highlighted three of these that were prominently seen in genes encoding developmental TFs. They are as follows: (1) promoter-region hypermethylation associated with silenced expression but needed only in a few of the cell types exhibiting such silencing; (2) DNA hypermethylation associated with down-modulated gene expression and suppressed or down-regulated enhancer activity; and (3) DNA hypermethylation upstream of promoters correlated with upregulation of gene expression probably by serving as a promoter border. Detailed examination of the relationships between various epigenetic profiles and the corresponding transcriptomes as well as the roles of the TFs in differentiation yielded insights into why a given gene region displays hypermethylated DMRs in certain cell types or tissues. Our study also offers paradigms for a better understanding of the contribution of abnormal DNA methylation to disease, e.g., cancer, the ICF syndrome, Duchenne muscular dystrophy, and congenital heart malformations, and for identifying genomic subregions where disease-related epigenetic aberrations are more likely to affect gene function.

## Methods

### Bioinformatics

Databases from the ENCODE and RoadMap projects ^23, 77^ with epigenetic and RNA-seq profiles used in the figures are available at the UCSC Genome Browser ^34^. The RRBS profiles for 18 types of cell culture samples used to determine myogenic differential methylation were previously described ^28^; the cell cultures were untransformed cell strains except for the LCLs. For tissue methylomes, we used bisulfite-seq profiles ^23, 33^ from the Bisulfite Sequencing Data hub rather than RRBS profiles because the two available RRBS methylomes for SkM were from individuals of advanced age (71 and 83 y) unlike the main BS-seq SkM sample which was a mixture of tissues from a 3 y male and a 34 y male. ^23^ In addition, bisulfite-seq data (which is not available for Mb) gives much more coverage than RRBS. We noticed that the RRBS profiles of SkM often displayed lower DNA methylation at Mb+Mt DM sites compared to BS-seq profiles of SkM from the same sites, which is probably attributable to the effects of aging on DNA methylation^78^. When more than one SkM bisulfite-seq track is shown, the extra tracks were psoas muscle from a 30 y female and separate analyses of the above two male samples. The chromatin state segmentation (chromHMM, AuxilliaryHMM) was from a hub for the Roadmap Epigenomics Project with the color code for chromatin state segmentation slightly simplified from the original ^23^, as indicated in the figures. The same sample mixture of 3 y and 34 y male psoas muscle was used for chromatin state segmentation as for bisulfite-seq. From the ENCODE project ^77^ we used the following UCSC Genome Browser tracks: RNA-seq (for tissues; not strand-specific), Massachusetts Institute of Technology ^79^; Transcription Levels by Long RNA-seq for poly(A)^+^ whole-cell RNA by strand-specific analysis on > 200 nt poly(A)^+^ RNA (for various cell cultures), Cold Spring Harbor Laboratories and RNA Subcellular CAGE Localization, RIKEN Omics Science Center. For visualizing RNA-seq tracks in the UCSC Genome Browser in figures, the vertical viewing ranges were 0 to 30 for cultured cells and 0 to 2 for tissues, unless otherwise specified. For Supplementary Tables S1b-S4b, quantification of RNA-seq for tissues was from the GTex database RPKM median values from more than 100 samples for each tissue type^80^ and for cell cultures was FPKM values as previously described.^28^

### Statistical analysis

Counts for Mb hypermethylated sites overlapping with ESC bivalent chromatin and/or CG islands were cross-tabulated and visualized using a mosaic plot (Fig 2b), a graphical method in which the area of each region is proportional to its relative magnitude. A Chi-square test of independence was used to assess the magnitude of the association between these two factors.

### Quantification of 5hmC and 5mC

The Epimark assay (New England Biolabs ^28^), which involved incubation of the DNA samples with T4 phage β-glucosyltransferase to glucosylate only 5hmC residues followed by cleavage at CCGG sites by restriction endonucleases (MspI, HpaII, or no digestion) and quantitative PCR (six reactions per sample) was done as previously described ^36^. The PCR primer-pairs for the analyzed CCGG sites in or upstream of *LXN, EBF3*, and *SIM1* are given in Supplementary Table S5a.

## Acknowledgments

We would like to thank Drs. Donald Comb, Rich Roberts, William Jack and Clotilde Carlow at New England Biolabs Inc. for research support and encouragement. We also thank Melody Badoo and the Tulane Cancer Center for help with the Cufflinks analysis of the ENCODE RNA-seq data.

